# Antibacterial T6SS1 cluster identification and bioinformatics characterization of their putative effectors in Shiga toxin-producing *Escherichia coli*

**DOI:** 10.1101/2024.03.05.583533

**Authors:** Nahuel A. Riviere, María Carolina Casabonne, Libia Y. Smith, Wanderson Marques da Silva, Ángel A. Cataldi, Mariano Larzábal

## Abstract

Shiga toxing-producing *Escherichia coli* (STEC) O22:H8 strain is a serotype occasionally isolated in Argentinian cattle. Preliminary works showed that the cattle carrying STEC O22:H8 strains could not be experimentally colonized by EHEC O157:H7 strain. The type 6 secretion system (T6SS) is one of the most versatile virulence mechanisms involved in delivering effectors, particularly the T6SS1 translocate antibacterial toxins effectors during bacterial competition for niche-space. In this work, we could evidence the molecular bases of the success of STEC O22:H8 (154) strain during bacterial competition against EHEC O157:H7 strains. The genome sequence of STEC O22:H8 (154) allowed us to identify a complete T6SS1 cluster. In addition, we identify and characterized several putative T6SS1-antibacterial effectors encoded inside the T6SS1 clusters and in genomic pathogenic islands. Competition assays against EHEC O157:H7 strain confirmed the antibacterial activity of STEC O22:H8 (154) strain *in vitro*. Considering the absent of T6SS1 in STEC strains, we proposed the recent horizontal transfer acquisition and the most probably donor belong to the same *Escherichia coli* species. A safe STEC O22:H8 (154) *Δstx* would be used as a new strategy to fight STECs in bovine intestinal colonization, leading in a reduction in beef contamination and consequently HUS cases in humans.

## 1. Introduction

Shiga toxing-producing *Escherichia coli* (STEC) are foodborne intestinal pathogens that cause hemolytic uremic syndrome (HUS) in humans ^17^. The serotype most frequently associated with HUS is O157:H7, which corresponds to *enterohemorrhagic Escherichia coli* (EHEC) strains belonging to a subgroup of STEC ^29^. STEC, whose primary reservoir is cattle, interact with the host, environment and with wide repertoire of pathogenic and commensals microbiota bacteria in the intestinal niche. STEC O22:H8 is a serotype that occasionally circulates in Argentinian cattle but that is rarely associated with human illness ^6^. Preliminary works showed that cattle carrying STEC O22:H8 (154) strain could not be experimentally colonized or persist in the rectoanal junction bovine by EHEC O157:H7 strain ^24^. Most probably STEC O22:H8 (154) can disrupt the balance of bovine intestinal microecosystem, including interbacterial competition with other STECs to facilitate their own gut colonization. However, the mechanism by which STEC O22:H8 (154) strain compete for the niche is still unknown.

The type 6 secretion system 1 (T6SS1) is a versatile virulence mechanisms involved in effectors toxin delivery in Gram-negative bacteria ^32^. In turn, T6SS1 is linked with a interbacterial competition activity for niche-space in some *Escherichia coli* pathotypes ^25^. However, to date, no T6SS1 has ever been identified in STECs bacteria. T6SS1 locus encodes minimal *core* components of 13 proteins (TssB, C, K, L, M, A, F, G, J, E, Hcp, ClpV and VrgG) to assemble the nanomachine apparatus ^4^. Currently, a wide repertoire of T6SS1 effectors toxins have been characterized. These include different catalytic toxins activities associated with interbacterial competition, such as proteinases (hydrolysis cytoplasmic proteins), muramidases or glycoside (hydrolysis of bacterial cell wall) ^10,37^, nucleases and deaminases (degradation and mutagenesis of nucleic acids) ^19^ and lipases (degradation of membrane structures) ^30^. Unlike other secretion systems, T6SS effectors do not contain a secretion signal and are translocated through covalent or *via cargo* (non-covalent) associated with VgrG spike proteins. Most of the T6SS1 effectors are encoded downstream of a *vgrG* and frequently contiguous an immunity protein to neutralize the toxicity of antibacterial toxin effectors ^10,14^. In particular, Rhs are a class of giant effectors proteins associated with T6SS that are composed of rearrangement hotspot regions (RHS). Most of Rhs are encoded out of the T6SS cluster and presents a consensus sequence PxxxxDPxGL that divides the protein in N- and C-terminal regions. The N-terminal region presents a PAAR motif to interact to VrgG at the tip of the T6SS and the variable C-terminal domain presents catalytic activity associated with a toxicity on the target cells (Figure 1).

**Figure 1:**
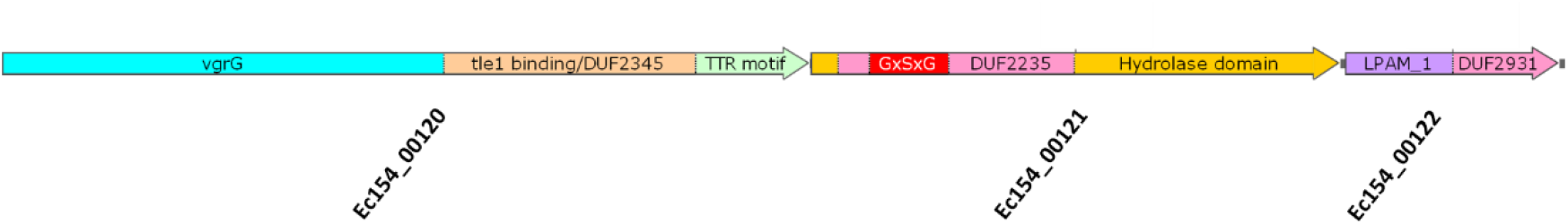
Rhs effectors proteins. These giant proteins composed of a consensus sequence PxxxxDPxGL that divides the protein into two regions, N- and C-terminal. The N-terminal sequence is composed of PAAR motif to interact to VrgG, rearrangement of hotspot (RHS) repeats-containing regions and a tyrosine/aspartate (YDxxGRL (I/T)) repeat region. The variable C-terminal domain presenting catalytic activity associated with a toxicity on the target cells.

In our study, we postulate the molecular bases of the success of STEC O22:H8 (154) strain during bacterial competition niche-space against EHEC O157:H7 strains. The genome sequence of STEC O22:H8 (154) allowed us to identify for the first time a T6SS1 cluster and several putative antibacterial effectors encoded inside and outside the cluster in STEC strains. In turn, we also postulate the recent horizontal transfer acquisition of T6SS1 by STEC strains. All that putative T6SS1 effector proteins, immunity proteins and adaptors were structurally characterized using bioinformatics studies as sequence database, 3D molecular modeling and domain analysis.

## 2. Materials and methods

### 2.1 Strains

STEC O22:H8 (154) (accession number CP067426-CP067431) and EHEC O157:H7 Rafaela II (accession number NZ_LAYW01000314.1) strains were taken from recto-anal mucosal swabs of healthy Holstein-Fresian calves in a dairies farms in Buenos Aires, Argentina ^1,24^. STEC O22:H8 (154) strain possesses two Shiga toxins (*stx1, stx2d*), *ehaA, ihA* and *lpfO113* genes involved in adhesion and biofilm formation ^24^. EHEC O157:H7 Rafaela II strain possesses two Shiga toxins (*stx2a* and *stx2c*), but is negative for *stx1*; in turn, the strain was positive for *eae* adhesin and *O157rbf* genes plasmid. EHEC O157:H7 Rafaela II also was classified as human hypervirulent clade 8 strain by lades according to an evaluation of a subset of 23 single nucleotide polymorphisms (SNPs) ^1^.

### 2.2 Bioinformatics analysis

The domain identification of putative T6SS1 effectors were performed using pfam (http://pfam.xfam.org/)11 and Batch Web CD Search-Tool (https://www.ncbi.nlm.nih.gov/Structure/bwrpsb/bwrpsb.cgi) ^23^. In addition, 3D molecular modeling assays and structure prediction studies to identify catalytic domains were performed using the PHYRE2 server (http://www.sbg.bio.ic.ac.uk/~phyre2/html/page.cgi?id=index) ^18^ and HHPRED (https://toolkit.tuebingen.mpg.de/tools/hhpred) ^39^, respectively. Proteins were analyzed by the Bastion6 server (https://bastion6.erc.monash.edu/) ^15^, which is an accurate predictor of potential T6SS effectors. The online server consists on a data-base of a wide genome prediction across 12 bacterial species, involving 54,212 protein sequences with more than 100 T6SS characterized effectors. BLASTp (https://blast.ncbi.nlm.nih.gov/Blast.cgi) was used to identify proteins homologous ^16^. All of these studies are listed in Supporting Information (Supplementary Table 1 and 2). The comparisons between the different T6SS1 or T6SS2 clusters was performed with the Artemis Comparison Tool (ACT) ^8^.

### 2.3 Phylogenetic analysis

Eighteen amino acid sequences of Tle1 protein from different *E. coli* pathotypes were retrieved from the National Center for Biotechnology Information database. In particular, more than 80 sequences of strains from a wide group of different serotypes were used. These included O157:H7 reference strains, non-O157 strains such as the “big six” group (O26, O45, O103, O111, O121, and O145) and O22:H8 among others. Non-intestinal *E. coli* strains such as EAEC, UPEC and ETEC were also used. All the sequences were aligned with the CLUSTAL W tool ^20^. The resulting alignment was used to carry out a Maximum-Likelihood phylogenetic inference using IQ-TREE v1.6.12 ^26^, with 10000 ultrafast bootstraps ^13^.

### 2.4 Competition assays

The study of the T6SS1^+^ STEC O22:H8 (154) strain antibacterial activity was performed by transforming a commensal and pathogenic T6SS1^-^prey strains, DH5-α and EHEC O157:H7 Rafaela II respectively, with pCRISPR-SacB-gDNA (Km^R^) plasmid to be selected by kanamycin. STEC O22:H8 (154) strain is sensitive to the kanamycin antibiotic. Overnight cultures of DH5-α, EHEC O157:H7 Rafaela II and STEC O22:H8 (154) were diluted to a final concentration of OD_600_ 0.5 in LB broth. The bacteria were centrifuged at low speed (6000 rpm) for 5 min and three washes were carried out with sterile PBS to eliminate the antibiotic residues. The T6SS1^+^ STEC O22:H8 (154) predator strain was mixed with T6SS1^-^DH5-α or EHEC O157:H7 Rafaela II prey strains in 0.5:1, 1:1 and 3:1 ratio. The mixtures were plated on 24-well plates containing LB agar. The plates were incubated for 16 h at 37°C. Then, the colonies were resuspended in 1 mL of sterile PBS. Serial dilutions were made and seeded on LB agar plates containing kanamycin (50 µg/mL). The assays were carried out in triplicate. Finally, the CFUs of surviving prey were quantified and the results were expressed as Log_10_ CFU/mL. Two-way ANOVA was performed to determine significance (p<0.0001).

## 3. Results

### 3.1 Antibacterial T6SS1 cluster identification in STEC O22:H8 (154) strain

The complete genome of STEC O22:H8 (154) strain was sequenced to identify the molecular basis of the colonization and interference competition against EHEC O157:H7 in the gut bovine niche ^24^. The analysis of the genome revealed the presence of genetic islands containing two completed T6SS clusters, which phylogenetically grouped with T6SS1 and T6SS2 according to the gene organization and cluster similarities (Figure 2A and B).

**Figure 2A and B:**
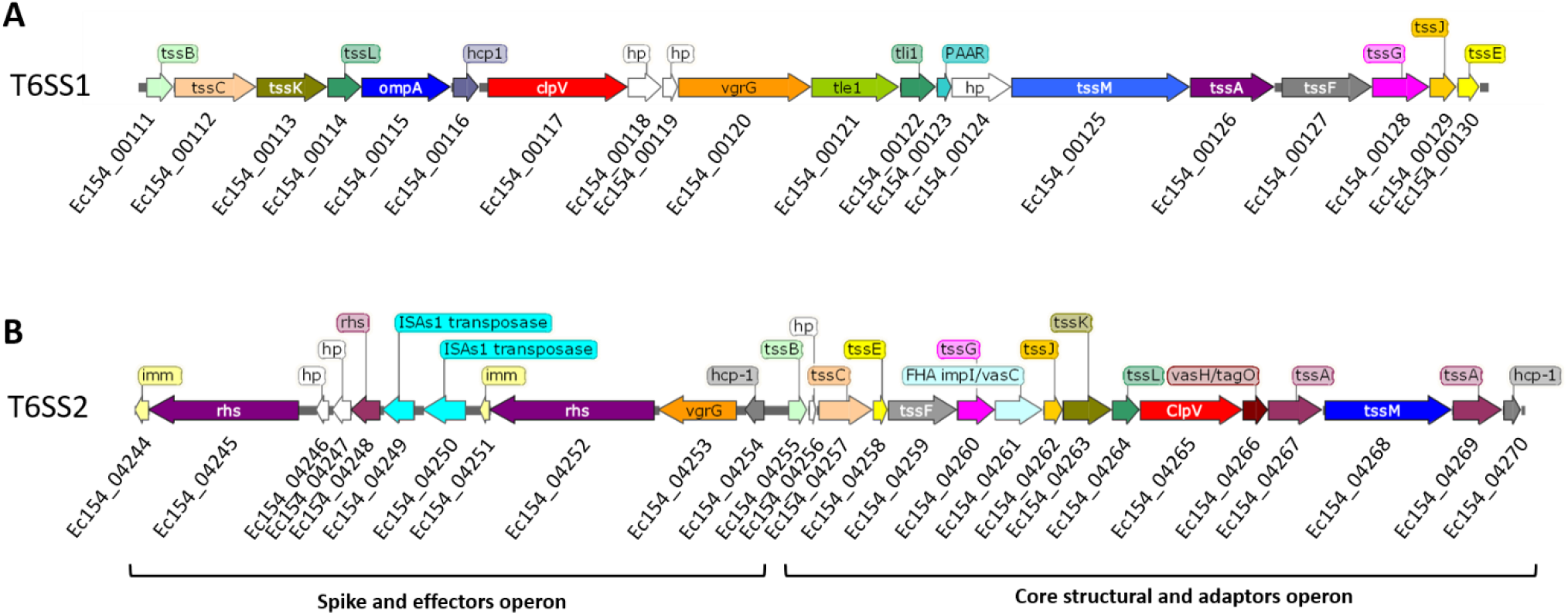
Diagram of T6SS1 and T6SS2 sequences in STEC O22:H8 (154) strain. Complete annotation and synteny genes in T6SS1 and T6SS2 clusters of STEC O22: H8 (154) strain. Both T6SS present all the essential components (core proteins) to develop the activity.

The T6SS1 *core* occupies the central region of a genetic island (Supplementary Figure 1). Upstream of the T6SS1 cluster, we found mostly proteins linked to RNA and DNA binding proteins, among which there are transcriptional regulators, mRNA inhibitors and cysteine deaminase proteins. Furthermore, we identified three putative T6SS immunity protein: *imm5* (EC154_00090), *tsi1* (EC154_00097) and *tsi3* (EC154_00103) but not the effectors associated. Downstream of the T6SS1 cluster, we found 8 transposases of IS3, IS66, IS600 and IS110 families. Other detected protein was a tyrosine-type recombinase/integrase (EC154_00146) with high homology to phage integrases.

### 3.2 Effectors characterization inside the T6SS1 STEC O22:H8 (154) cluster

*In silico* analysis of STEC O22:H8 (154) T6SS1 cluster allowed us to identify three potential genes associated with translocation and effector activities. The genes are encoded in tandem inside the locus (from Ec154_00120 to 22) (Figure 3). Ec154_00120 gene corresponds to putative VgrG1 spike. Ec154_00121, encoded downstream of the *vgrG1*, corresponds to putative type 6 secretion protein associated with hydrolase-toxin lipase effectors. According to a comparison analysis of the sequences, Ec154_00121 presented a 95.9% identity with T6SS effector phospholipase Tle1-EAEC 042. Tle1 is linked to an activity on the membrane of prokaryotic cells during interbacterial competition mediated by T6SS1. Tle1 represents a cargo-type translocation effector, since it needs to interact with VgrG1 to be translocated. The interaction occurs through TTR motif of VgrG1, which is directly associated with the N-terminal region of Tle1. This TTR motif is conserved in *vgrG* (Ec154_00120), which would allow Tle1 to bind at the T6SS tip complex. Usually, the effector-immunity complexes are encoded in tandem. The self-protection of Tle1 toxin is an outer membrane lipoprotein, named Tli1 that binds to Tle1 and thus inhibits the phospholipase activity. Ec154_00122 gen presents 97.8% identity to Tli1 immunity protein of EAEC 042 and has the two characteristic domains in the N-terminal region: DUF2931 and a lipoprotein attachment LPAM_1.

**Figure 3:**
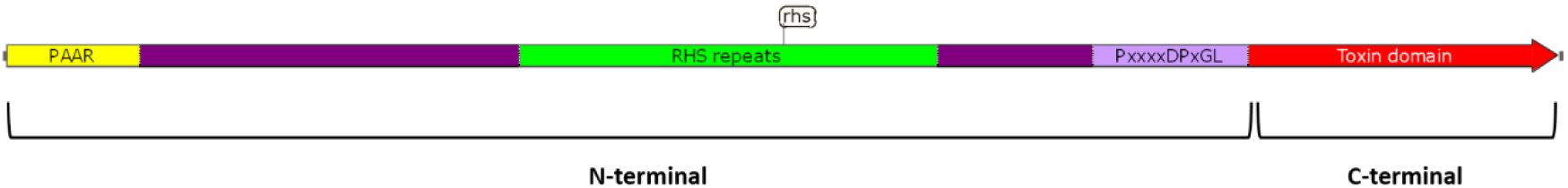
Effectors inside the T6SS1 cluster. The T6SS1 cluster presents a VgrG1 with Tle1 Binding/DUF2345 (orange) domain and TTR (*transthyretin-like domain*) motif (green) in the C-terminal region. Tle1 also presents GxSxG motif (red) inside the DUF2235 domain (pink) associated with *α/β* hydrolase domain. GxSxG motif is usually found in phospholipases of Tle1-4 family. All the sequences present a hydrolase-toxin lipase effector (yellow). Downstream of Tle1, there is the self-*protection lipoprotein Tli1 that* has a lipoprotein attachment LPAM_1 characteristic domain in the N-terminal region (violet) and a DUF2931 (pink) domain in the C-terminal region.

### 3.3 Effectors characterization outside of T6SS1 STEC O22:H8 154 cluster

*In silico* analysis of the STEC O22:H8 (154) strain genome revealed six pathogenic islands with potential effectors linked to T6SS1.

#### 3.3.1 Pathogenic island 1

This pathogenic island presents an operon composed of *vgrG1* (Ec154_03857), *eagR* (Ec154_03858) and three *rhs* genes with their respective tandem-encoded immunity proteins (Ec154_03859/60, Ec154_03862/3 and Ec154_03866/7) (Figure 4A). VgrG1 (Ec154_03857) presents a 98.9% identity with VgrG_O1_ HMPREF9348_01321 from *E. coli* MS 145-7 and 98.3% identity with VgrG_O1_ ECDEC10E_0686 form *E. coli* O26:H11 ECDEC10E and VgrG_O1_ AAF13_16105 STEC004. The *eagR* (***e****ffector-****a****ssociated* ***g****ene with* ***R****hs*) gene is encoded downstream of *vgrG*, it has a DUF1795/DcrB domains and belongs to the family of conserved accessory proteins. EagR has a key role for the T6SS1-dependent delivery of toxic Rhs-effectors acting as a professional chaperones, and is typically encoded between *vgrG* and *rhs* genes ^19^. This type of chaperone is normally found associated with effectors linked to antibacterial activity.

**Figure 4A, B, C and D:**
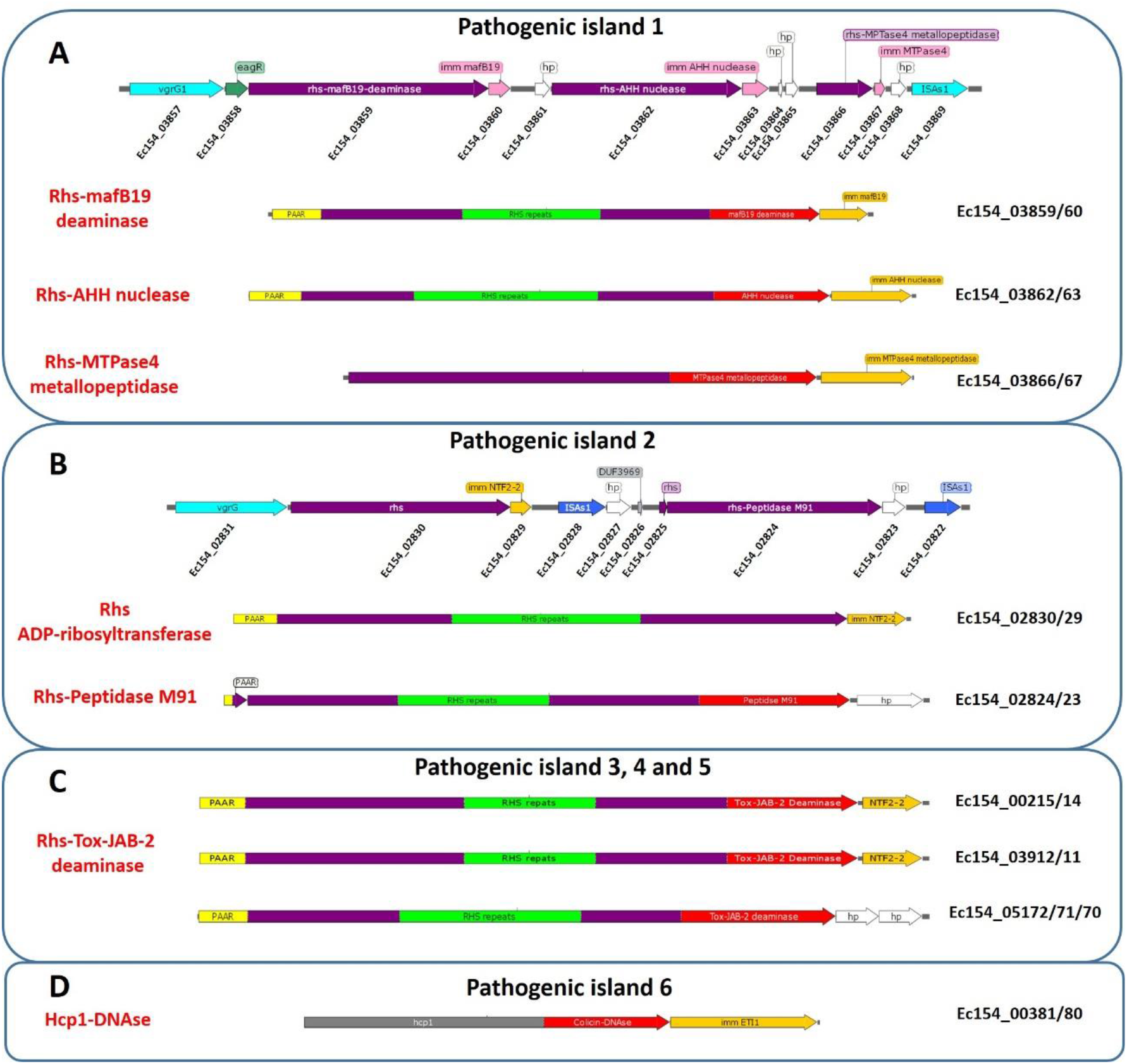
Diagram of pathogenic islands: The sequence of the different potential effectors is represented. VgrG (light blue), Hcp protein (gray), Rhs family proteins (violet) with their respective immunity proteins downstream (orange) and hypothetical proteins (white). Within the sequence of the Rhs proteins we can identify rearrangement hotspot (RHS) repeat-containing regions (green), the presence of the PAAR motif to interact with VrgG in the N-terminal region (yellow) and the catalytic toxin C-terminal domain (red).

Ec154_03859 gene is encoded downstream of *vgrG* and *eagR*, thus suggesting a potential effector activity linked to T6SS. Ec154_03859 possesses typical characteristics of RhsA protein, as evidenced by the presence of PAAR motif and RHS repeats. In the C-terminal region, it presents a mafB19-nucleotic deaminase domain and the HxT motif related to novel family toxin proteins involved in interbacterial competition ^22^. This domain has 100% identity in relation to other mafB19-nucleotic deaminase domains of T6SS1 effector already characterized as HMPREF9348_01319 *E. coli* MS 145-7 and EFO59306 from *E. coli* strains ^22^. The total identity between these proteins is 97.8%. The mafB19-nucleotic deaminase domain is related to nucleic acid editing, modifications and defensive deployment leading to DNA hypermutation in the cell target; the nucleoacidic substrate of the Rhs-mafB19 effectors, however, remains elusive ^22^. Downstream of Rhs-mafB19, we identified its antagonist immunity protein (Ec154_03860), which presents 100% identity with HMPREF9348_01318 mafB19-nucletic deaminase immunity protein from *E. coli* MS 145-7 strain. This finding suggest its inhibitory function.

Continuous downstream the operon, we identified a predicted RhsA (Ec154_03862) with a C-terminal containing nuclease AHH (*alanine-histidine-histidine*) motif. The AHH motif shows a 98% identity with T6SS1 Rhs-AHH effectors already characterized, such as KIH17755 from *E. coli* CVM N33653PS and ECDEC10E_0684 from *E. coli* O26:H11 DEC10E. In the AHH motif, the first histidine forms one of the catalytic metal-chelating ligands, whereas the second histidine contributes to the active site directing the water for phosphoester hydrolysis ^38^. The Rhs-AHH effectors are the most dominant nuclease found in 66 bacterial species associated to T6SS ^22^. Moreover, researchers have reported many Rhs-AHH in *E. coli* with T6SS1 antibacterial activity ^33^. Ec154_03863 protein, which is encoded downstream of Rhs-AHH, has 99.4% identity with ECDEC10E_0683 Rhs-AHH nuclease immunity protein of *E. coli* O26:H11 DEC10E.

Close to Ec154_03863 immunity protein, we identified another RhsA protein (Ec154_03866) with a zincin-like metallopeptidase-4 domain (Tox-MPTase4) C-terminal region. This domain displays a 98.2% identity with Rhs-MPTase4 effector proteins of T6SS1, such as AAF13_16115 from *E. coli* O104:H4 C227-12 and AOM81335 from STEC004. The MPTase4 domain exhibits activity against membrane and wall in target cells ^22^. However, *rhsA* Ec154_03866 sequence lacks the first 1150 N-terminal amino acids and the central region, in which PAAR and RHS repeat motifs are commonly encoded. Tox-MPTase4 immunity protein (Ec154_03867) is encoded downstream of the effector protein and presents a high identity (98.6%) with other cognate immunity proteins already characterized as AAF13_16120 from *E. coli* O104:H4 C227-11 and *cti1* of STEC004 strains.

#### 3.4.2 Pathogenic island 2

The pathogenicity island 2 encodes VgrG (Ec154_02831), two putative Rhs effectors (Ec154_02830 and Ec154_02824) and the respective cognate immunity proteins (Ec154_02829 and Ec154_02823) associated with T6SS (Figure 4B). This island lacks adapter proteins or chaperones. Structural analyzes of Rhs Ec154_02830 by Phyre2 revealed that the C-terminal region possesses a low confidence with an ADP-ribosyltransferase (ART) family toxin domain.

Close to Ec154_02830 gene is encoded Ec154_02824, a predicted RhsA with a zinc metallopeptidase M91 domain at the extreme C-terminus associated with a T6SS1 effector activity. The metal-binding domain consists of a HExxH consensus motif in the C-terminal of Ec154_02824. The two histidine residue ligands, the catalytic Zn2+ and the glutamic acid residue are involved in the nucleophilic peptidase activity ^3^. RhsA Ec154_02824 has no PAAR motif, because of a premature stop codon that divided the protein in Ec154_02824 and Ec154_02825 genes. On the other hand, no homology or identity with immunity proteins could be determined for the hypothetical protein Ec154_02823 encoded downstream of RhsA.

#### 3.4.3 Pathogenic island 3, 4 and 5

These islands encode three putative effector proteins Ec154_00215, Ec154_05172 and Ec154_03912, which were significantly predicted as Rhs proteins (Figure 4C). In turn, each of these proteins has in the C-terminal region DUF4329 domains with homology to Rhs-Tox-JAB-2 potentially associated with RNase activity ^22^. The cognate immunity protein NTF2 family 2 Ec154_00214 and Ec154_03911 are encoded downstream of Rhs-Tox-JAB-2 Ec154_00215 and Ec154_03912, respectively. The island that contains Rhs-Tox-JAB-2 Ec154_05172 does not code for the immunity protein NTF2-2. However, the RNase activity of Rhs-Tox-JAB-2 Ec154_05172 could be blocked by Ec154_00214 or Ec154_03911 NTF2-2 proteins.

#### 3.4.4 Pathogenic island 6

An *hcp* (Ec154_00381) gene is encoded outside of the T6SS cluster or the *vgrG* islands. This orphan Hcp protein has a HNH endonuclease and colicin E7/E9 endonuclease domains in the C-terminal region (Figure 4D). The Hcp with C-terminal Extension Toxins are designated as Hcp-ET and are proteins widely present in the *Enterobacteriaceae*. Hcp-ETs fulfill both the structural function in the formation of the Hcp tail tube as well as an effector function in the target cell ^5,14^.Hcp (Ec154_00381) shows a 99% identity with Hcp-ET1 from *E. coli* O104:H4 C227-11 and *E. coli* 55989 strains, which presents a nonspecific cleavage HNH-DNAse activity. Most likely, Hcp forms hetero-hexameric rings with Hcp-ETs and the catalytic HNH-DNAse domain is oriented inside the tube. The ETI1 immunity protein neutralized this DNase activity toxin. Ec154_00380, encoded downstream of Hcp-ET1, shows 99.4% identity with ETI1 of *E. coli* O104:H4 C227-11 and *E. coli* 55989 strains; which indicates that the toxin is also encoded in tandem in STEC O22:H8 (154) with its immunity protein.

### 3.3 Bioinformatic identification of T6SS2 clusters in STEC O22:H8 154 strain

According to bioinformatic analysis, the STEC O22:H8 (154) strain contains a complete T6SS2 island divided into two operons (Figure 2B). One of the operons contains the *core* structural and adaptors genes, whereas the other presents the components of the spike, effectors and immunity proteins. ACT comparative analysis of T6SS2 sequences showed the same operon distribution and high synteny between STEC O22:H8 (154) and EHEC O157:H7 EDL933 reference strain (Supplementary Figure 2). Particularly, T6SS2 is widely distributed in STECs bacteria and has been shown by comparative analysis to exhibit no significance difference in the operon distribution and synteny between different strains ^35^. Recently, T6SS2 of STEC strains has been associated with intramacrophage survival and not with interbacterial competence ^36^. Macrophages phagocytose STEC strains adhering to the intestinal epithelium and induce the production and release of reactive oxygen species (ROS) to eliminate the bacteria. In the face of the oxidative response, T6SS2 translocates effectors with catalase activity to the macrophage cytosol reducing intracellular ROS levels and allowing STEC survival and replication inside the macrophage ^35,36^.

### 3.6 T6SS clusters acquisition by STEC O22:H8 (154) strain

Previous research has suggested that the horizontal transfer of T6SS clusters in *E. coli* has not been recently acquired, since GC content frequency of clusters is quite similar to that of the chromosome ^9^. We observed that STEC O22:H8 (154) has a 50.83% GC chromosome content, which is similar to T6SS1 (50.74%) and T6SS2 (52.57%) clusters. Nevertheless, these islands are located close to tRNA and contain transposase and other mobile elements. This finding strongly suggests acquisition by lateral transfer from other bacteria whose genome has a similar GC% composition. Particularly, T6SS2 is inserted close to tRNA Asp, inside the *yaf* operon (Supplementary Figure 3A). T6SS2 location is conserved in most of STEC including EHEC O157:H7 strain, suggesting that the transfer event has not occur recently. On the other hand, T6SS1 is inserted in STEC O22:H8 (154) close to tRNA SelC inside of *yicIJH* operon, which is involved in sugars efflux ^28^ (Supplementary Figure 3B). The *yicIJH* operon is part of the common genetic backbone in most of the *E. coli*, including symbionts and pathogenic strains ^27^. We could even observe that STEC O22:H8 RM10809-C3, a strain isolated and sequenced by Qiu Carter et al, has the same tRNA SelC organization locus, but without T6SS1 insertion island ^7^.

The presence of T6SS1 has never been described in STEC strains, so an exhaustive search for the cluster was performed in a wide repertoire of sequences of different STEC serotypes in Genbank. This data was used to construct a phylogenetic tree based on the conservation of the Tle1 effector encoded within T6SS1 (Figure 5). The presence of the T6SS1 was detected only in 5% of the STEC strains evaluated, unlike T6SS2 that was present in 42% of the STEC strains. Evolutionary analysis showed that the T6SS1 of STEC O22:H8 (154) is clustered with two strains of serotypes O91:H14 and O76:H19 isolated from human stool, and MFDS1006657 and STEC639 strains of undetermined serotypes from beef ribs and sheep isolates respectively. T6SS1 is highly prevalent in EAEC strains, however, we observed that it diverges from T6SS1 of STEC O22:H8 (154) and is closer to the neighbourhood of T6SS1 of ETEC and UPEC.

**Figure 5:**
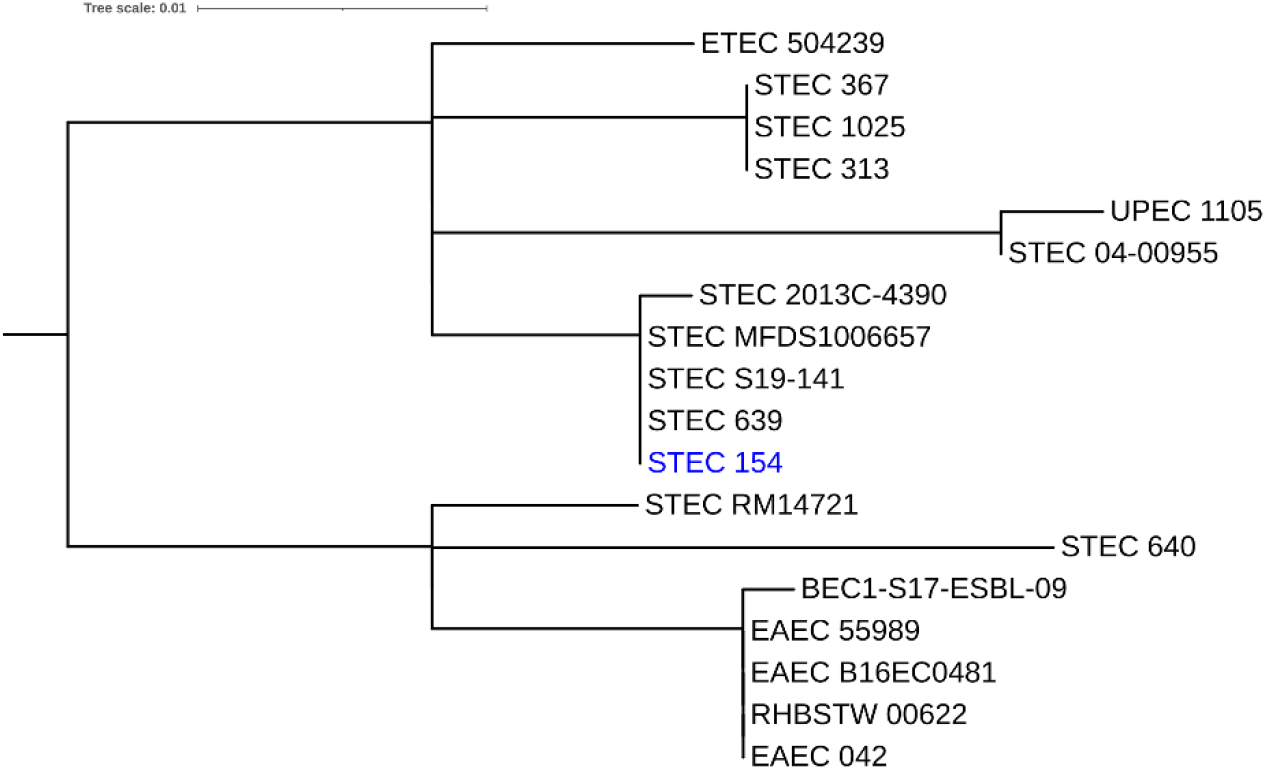
Phylogenetic tree based on Tle1 of T6SS1: Sequence from different *E. coli* pathotypes were aligned with CLUSTAL W tool to construct a phylogenetic tree.

### 3.7 Competition assays predator-prey

Intraspecies competition assays were performed to demonstrate the potential antibacterial role of T6SS1 of STEC O22:H8 (154). We co-cultured STEC O22:H8 (154) in the presence of two commensal and pathogenic T6SS1 negative *E. coli* preys: DH5-α or EHEC O157:H7 Rafaela II strains respectively. In presence of STEC O22:H8 (154) in a ratio of 0.5:1, DH5-α reduced its survival in 3 orders of magnitude, whereas EHEC O157:H7 Rafaela II was reduced in 1 order. The increase of predator concentration strain to 3:1 ratio caused higher reductions for DH5-α (4 orders) and EHEC O157:H7 Rafaela II (3 orders) strains respectively (Figure 6). Similar analyzes using the STEC O22:H8 (154) strain filtered supernatant demonstrated the absence of secretory proteins with bactericidal activities against DH5-α or EHEC O157:H7 Rafaela II strains. Competition assays between EHEC O157:H7 Rafaela II (T6SS2^+^) vs DH5-α strains showed no reduction in the survival of DH5-α; which confirms that T6SS2 of STEC strains has no antibacterial activity (data not shown).

**Figure 6:**
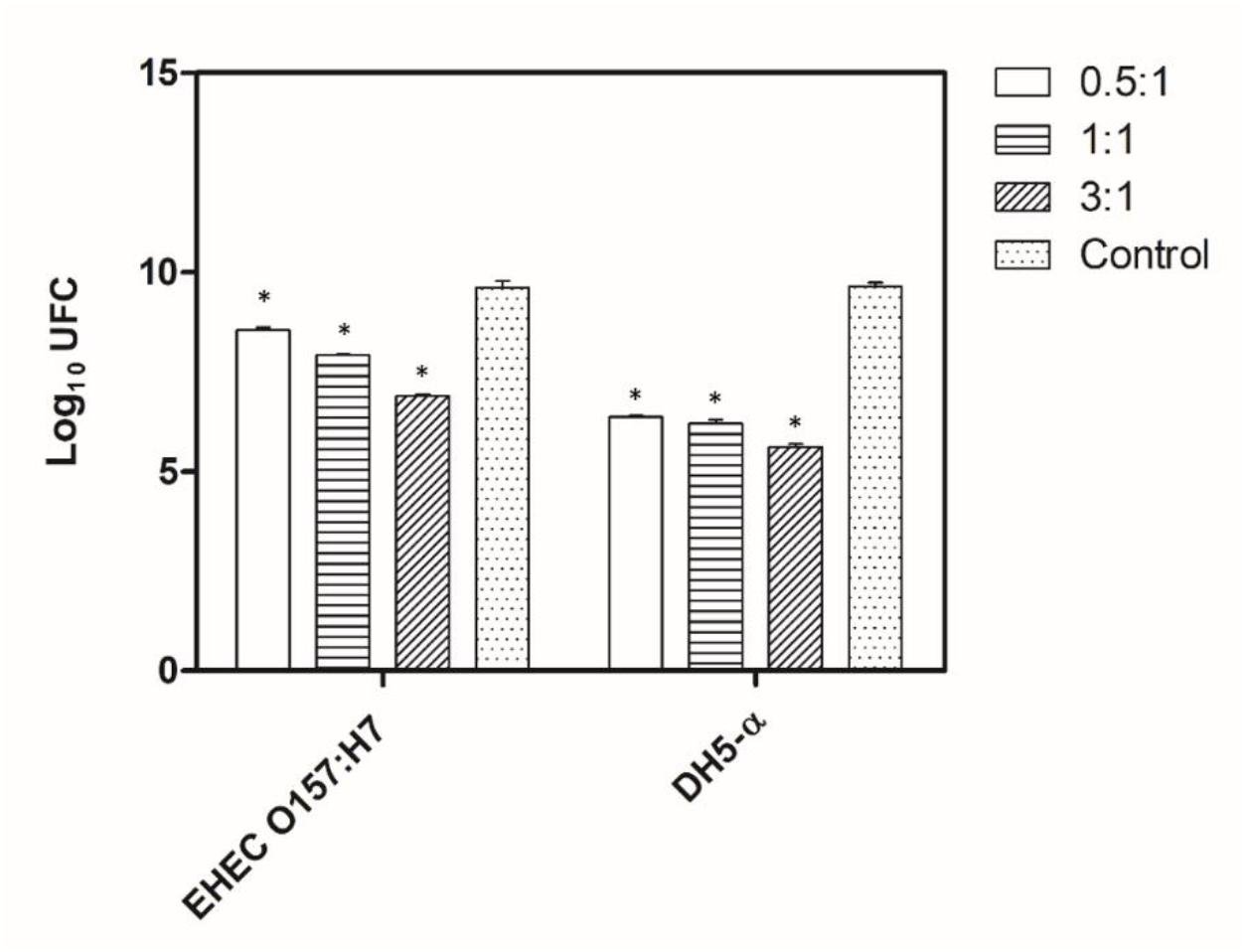
Competition assays. Co-cultures of STEC O22:H8 (154) strain in the presence of *E. coli* DH5-α or EHEC O157:H7 Rafaela II strains. The survival of prey bacteria was quantified by log_10_ UFC/ml. The ratio of STEC O22:H8 (154) vs *E. coli* DH5-α or EHEC O157:H7 Rafaela II were represented by 0.5:1, 1:1 and 3:1. Two-way ANOVA was performed as a statistical analysis. The asterisk represents a significant p<0.0001 value.

## 4. Discussion

T6SS1 effectors can be encoded in or out of the cluster and be translocated through covalent or *via cargo*. The best-characterized effector of O22:H8 (154) strain was Tle1, encoded inside the cluster that preserves the catalytic triad composed of Ser-197, Asp-245 and His-310 amino acids associated with phospholipase A1 and A2 activities. Flaugnatti et al. confirmed that a substitution of Ser-197 by an alanine residue abolished a phospholipase activity in Tle1-EAEC 042 protein ^12^. Tle1 also presents GxSxG motif, which is usually found in phospholipases of Tle1-4 family at 195-199 position ^31^. Considering the conservation of the catalytic triad and the GxSxG motif, Tle1 of O22:H8 (154) could be associated with TTR motif of VgrG and be translocated *via cargo* into the periplasm degrading the cell membrane of the prey cell. Flaugnatti et al. also demonstrated that Tle1 activity was completely abolished in the presence of Tli1 anchored to the membrane ^12^. Therefore, the conservation of the domains in the N-terminal region of Tli1 from O22:H8 (154) would indicate that it could have a self-protective activity against Tle1.

An abundance of potential Rhs effectors were identified encoded in pathogenicity islands (PIs) outside the T6SS1 cluster. *rhs* genes are widely distributed in *E. coli* and are linked by horizontal gene transfer T6SS effectors acquisitions. All potential encoded Rhs effectors presented rearrangement hotspot (RHS) repeats-containing regions and a consensus sequence PxxxxDPxGL. However, not all of them presented PAAR motifs in the N-terminal region. Ec154_02824 (peptidase M91) and Ec154_03866 (MTPase4) are two examples of Rhs that lost the PAAR motif. In case of Ec154_02824 the loss of the motif is due to a premature stop codon divided the sequence in Ec154_02824 and Ec154_02825 genes. The inability to associate with VgrG by the PAAR motif would prevent translocation of the Rhs effector *via cargo* by the T6SS1.

Rhs-mafB19-nucleotide deaminase, Rhs-AHH and Rhs-MTPase4 are encoded downstream of VgrG and EagR chaperone in PI1, indicating that they could be associated effectors. Ma et al. have demonstrated that the delivery of Rhs-mafB19-nucleotic deaminase, Rhs-AHH and Rhs-MTPase4 effector toxins are dependent on the VgrGO1 and EagR chaperone activity ^22^. However, they did not observe the presence of these effectors on the same island. In addition, VgrG_O1_ which presents a complete homology with VgrG (Ec154_03857), is involved in translocation *via cargo* and participates in the antibacterial activity induced by T6SS1 ^22^. The association between VgrG_O1_ and Rhs-mafB19 or AHH would occur through the EagR chaperone thus allowing the translocation of the effectors to the target cell. Thus, different Rhs effector proteins encoded in the operon could use the same VgrG and the EagR chaperone. This operon organization could confer to STEC O22:H8 (154) better versatility and efficiency to translocate effectors proteins by T6SS1.

Rhs ADP-ribosyltransferase and Rhs-Peptidase M91 are encoded downstream of VgrG in PI2. Unlike PI1, PI2 has no EagR helper protein. Recently, Ting et al. have characterized a Rhs effector protein named Tre1 with ADP-ribosyltransferase interbacterial activity exported by the T6SS in *Serratia proteamaculans* ^34^. Tre1 modifies the essential bacterial tubulin-like protein (FtsZ) and blocks its capacity to polymerize affecting the bacterial structure. The authors did not determine the need for a chaperone protein to be translocated to the target bacterium. There is no evidence in *E. coli* of Rhs effectors that is inhibit the polymerization activity. On the other hand, Rhs-Peptidase M91 presents a domain associated with the M91 zinc metallopeptidase family because of the presence of an HEXXH motif. The domain has homology with the characterized effector NleD of enteropathogenic *Escherichia coli* (EPEC) that is translocated to epithelial cells by the type 3 secretion system. NleD is a virulence factor that helps to inactivate the inflammatory response by cleavage of host mitogen-activated protein kinase 9 ^2^. So far, no T6SS-associated Rhs-Peptidase M91 proteins have been identified, so its potential endopeptidase activity in the cytoplasm of the target bacteria, as well as its substrate, is unknown. Both Rhs potential effectors are an important object of study.

Inside the PI3, 4 and 5 identified three orphans Rhs-Tox-JAB-2 encoded without VgrG or helpers proteins. The potentially catalytic activity of Tox-JAB-2 is associated with RNase ^22^. However, although it is clear that Rhs-Tox-JAB-2 has T6SS effector characteristics, its catalytic activity and substrate is still unknown. Its repeats could be due to independent acquisitions or duplications in the genome of STEC O22:H8 (154) strain.

Finally, Hcp encoded in PI6, exhibits characteristics of Hcp-ET1 with catalytic HNH-DNAse domain. According to Ma et al.’s study, Hcp-ET1 has a cytotoxicity activity by degrading target DNA of recipient during the antibacterial competition ^21^. Furthermore, T6SS2, but not T6SS1, is responsible for the delivery of Hcp-ET1 to the cytoplasm of the target cells ^21^.

## 5. Conclusion

Martorelli et al. have shown that STEC O22:H8 (154) strain might compete with EHEC O157:H7 strain at the bovine recto-anal junction, making STEC O22:H8 (154) carrying-calves less susceptible to EHEC O157:H7 strain colonization and reducing the excretion of bacteria to the environment. The study of the genome sequence of STEC O22:H8 (154) allows us to support that T6SS1 is involved in bacterial competence *in vitro* and *in vivo* against EHEC O157:H7. The presence of two complete T6SS operons and several pathogenic island containing putative effectors probably confers a fitness advantage to STEC O22:H8 (154) during the pathogenesis. Once the bacteria colonize the bovine gut, their persistence deepens on the T6SS2 and its role against the host innate immune response. Then, T6SS1 would allow the bacteria to compete for the environmental niches against rival bacteria. However, further experimental supports are needed to understand the involvement of T6SS1 and Rhs effectors during pathogenesis and bacterial competition.

Some *E. coli* species have T6SS1 clusters but, interestingly, to date no T6SS1^+^ STEC strains had been described. In view of the low proportion of STEC strains with T6SS1, the presence of several integrase families within the T6SS1 island and its close insertion into a tRNA region suggest recent horizontal transfer. Most likely this acquisition event involving another *E. coli* species as a donor.

STEC O22:H8 is a serotype occasionally isolated in Argentinian cattle that contains *stx1* and *stx2d*, although it is rarely associated with human illness. Through genic edition, a safe STEC O22:H8 (154) *Δstx* strain could be obtained to use as a potential bovine probiotic. As already shown, STEC O22:H8 (154) efficiently colonizes and persists in the gut bovine, the same niche that shares with EHEC O157:H7 strain, the serotype responsible for worldwide outbreaks of HUS. In this way, experimentally inoculated bovines would avoid EHEC O157:H7 intestinal colonization and potentially would block the colonization of other STEC T6SS1 negative of the Big Six (non-O157) serotypes groups. STEC O22:H8 (154) *Δstx* would be used as a new strategy to fight STEC intestinal colonization, which would lead to a reduction in beef contamination and consequently HUS cases in humans.

## References

1. Amigo N, Mercado E, Bentancor A, Singh P, Vilte D, Gerhardt E, Zotta E, Ibarra C, Manning SD, Larzábal M, Cataldi A. Clade 8 and clade 6 strains of Escherichia coli O157:H7 from cattle in Argentina have hypervirulent-like phenotypes. PLoS ONE. 2015 [accessed 2020 Nov 6];10(6). https://pubmed.ncbi.nlm.nih.gov/26030198/. doi:10.1371/journal.pone.0127710

2. Baruch K, Gur-Arie L, Nadler C, Koby S, Yerushalmi G, Ben-Neriah Y, Yogev O, Shaulian E, Guttman C, Zarivach R, Rosenshine I. Metalloprotease type III effectors that specifically cleave JNK and NF-κB. EMBO Journal. 2011;30(1):221–231. 10.1038/emboj.2010.297. doi:10.1038/emboj.2010.297

3. Berni B, Soscia C, Djermoun S, Ize B, Bleves S. A type VI secretion system trans-kingdom effector is required for the delivery of a novel antibacterial toxin in Pseudomonas aeruginosa. Frontiers in Microbiology. 2019 [accessed 2020 Nov 6];10(JUN). https://pubmed.ncbi.nlm.nih.gov/31231326/. doi:10.3389/fmicb.2019.01218

4. Boyer F, Fichant G, Berthod J, Vandenbrouck Y, Attree I. Dissecting the bacterial type VI secretion system by a genome wide in silico analysis: What can be learned from available microbial genomic resources? BMC Genomics. 2009 [accessed 2020 Nov 6];10. https://pubmed.ncbi.nlm.nih.gov/19284603/. doi:10.1186/1471-2164-10-104

5. Brunet YR, Hénin J, Celia H, Cascales E. Type VI secretion and bacteriophage tail tubes share a common assembly pathway. EMBO Reports. 2014 [accessed 2020 Nov 13];15(3):315–321. /pmc/articles/PMC3989698/?report=abstract. doi:10.1002/embr.201337936

6. Cadona JS, Bustamante A V., González J, Sanso AM. Genetic relatedness and novel sequence types of non-O157 Shiga toxin-producing Escherichia coli Strains Isolated in Argentina. Frontiers in Cellular and Infection Microbiology. 2016 [accessed 2020 Nov 6];6(AUG):93. https://www.frontiersin.org. doi:10.3389/fcimb.2016.00093

7. Carter MQ, Pham A, He X, Hnasko R. Genomic Insight into Natural Inactivation of Shiga Toxin 2 Production in an Environmental Escherichia coli Strain Producing Shiga Toxin 1. Foodborne Pathogens and Disease. 2020 [accessed 2020 Nov 6];17(9):555–567. https://www.liebertpub.com/doi/10.1089/fpd.2019.2767. doi:10.1089/fpd.2019.2767

8. Carver TJ, Rutherford KM, Berriman M, Rajandream MA, Barrell BG, Parkhill J. ACT: The Artemis comparison tool. Bioinformatics. 2005 [accessed 2020 Nov 23];21(16):3422–3423. https://www.webact.org. doi:10.1093/bioinformatics/bti553

9. Cascales E. The type VI secretion toolkit. EMBO reports. 2008 [accessed 2020 Nov 6];9(8):735–741. https://onlinelibrary.wiley.com/doi/abs/10.1038/embor.2008.131. doi:10.1038/embor.2008.131

10. Dong TG, Ho BT, Yoder-Himes DR, Mekalanos JJ. Identification of T6SS-dependent effector and immunity proteins by Tn-seq in Vibrio cholerae. Proceedings of the National Academy of Sciences of the United States of America. 2013 [accessed 2020 Nov 6];110(7):2623–2628. https://www.pnas.org/cgi/doi/10.1073/pnas.1222783110. doi:10.1073/pnas.1222783110

11. El-Gebali S, Mistry J, Bateman A, Eddy SR, Luciani A, Potter SC, Qureshi M, Richardson LJ, Salazar GA, Smart A, Sonnhammer ELL, Hirsh L, Paladin L, Piovesan D, Tosatto SCE, Finn RD. The Pfam protein families database in 2019. Nucleic Acids Research. 2019 [accessed 2020 Nov 23];47(D1):D427–D432. https://pfam.xfam.org. doi:10.1093/nar/gky995

12. Flaugnatti N, Rapisarda C, Rey M, Beauvois SG, Nguyen VA, Canaan S, Durand E, Chamot-Rooke J, Cascales E, Fronzes R, Journet L. Structural basis for loading and inhibition of a bacterial T6 SS phospholipase effector by the VgrG spike . The EMBO Journal. 2020;39(11):1–14. doi:10.15252/embj.2019104129

13. Hoang DT, Chernomor O, Von Haeseler A, Minh BQ, Vinh LS. UFBoot2: Improving the Ultrafast Bootstrap Approximation. Molecular biology and evolution. 2018 [accessed 2022 Mar 28];35(2):518–522. https://pubmed.ncbi.nlm.nih.gov/29077904/. doi:10.1093/MOLBEV/MSX281

14. Hood RD, Singh P, Hsu FS, Güvener T, Carl MA, Trinidad RRS, Silverman JM, Ohlson BB, Hicks KG, Plemel RL, Li M, Schwarz S, Wang WY, Merz AJ, Goodlett DR, Mougous JD. A Type VI Secretion System of Pseudomonas aeruginosa Targets a Toxin to Bacteria. Cell Host and Microbe. 2010 [accessed 2020 Nov 6];7(1):25–37. /pmc/articles/PMC2831478/?report=abstract. doi:10.1016/j.chom.2009.12.007

15. Jiaweiwang J, Yang B, Leier A, Marquez-Lago TT, Hayashida M, Rocker A, Zhang Y, Akutsu T, Chou KC, Strugnell RA, Song J, Lithgow T. Bastion6: A bioinformatics approach for accurate prediction of type VI secreted effectors. Bioinformatics. 2018 [accessed 2020 Nov 23];34(15):2546–2555. https://academic.oup.com/bioinformatics/article/34/15/2546/4934942. doi:10.1093/bioinformatics/bty155

16. Johnson M, Zaretskaya I, Raytselis Y, Merezhuk Y, McGinnis S, Madden TL. NCBI BLAST: a better web interface. Nucleic acids research. 2008 [accessed 2020 Nov 23];36(Web Server issue). https://pubmed.ncbi.nlm.nih.gov/18440982/. doi:10.1093/nar/gkn201

17. Kaper JB, Nataro JP, Mobley HLT. Pathogenic Escherichia coli. Nature Reviews Microbiology. 2004 [accessed 2020 Nov 6];2(2):123–140. https://pubmed.ncbi.nlm.nih.gov/15040260/. doi:10.1038/nrmicro818

18. Kelley LA, Mezulis S, Yates CM, Wass MN, Sternberg MJ. Trabajo práctico No 13 . Varianzas en función de variable independiente categórica. Nature Protocols. 2016;10(6):845–858. 10.1038/nprot.2015-053. doi:10.1038/nprot.2015-053

19. Koskiniemi S, Lamoureux JG, Nikolakakis KC, De Roodenbeke Ctk, Kaplan MD, Low DA, Hayes CS. Rhs proteins from diverse bacteria mediate intercellular competition. Proceedings of the National Academy of Sciences of the United States of America. 2013 [accessed 2020 Nov 6];110(17):7032–7037. https://www.pnas.org/content/110/17/7032. doi:10.1073/pnas.1300627110

20. Larkin MA, Blackshields G, Brown NP, Chenna R, Mcgettigan PA, McWilliam H, Larkin MA, Blackshields G, Brown NP, Chenna R, McGettigan PA, McWilliam H, Valentin F, Wallace IM, Wilm A, Lopez R, Thompson JD, Gibson TJ, Higgins DG. Clustal W and Clustal X version 2.0. Bioinformatics (Oxford, England). 2007 [accessed 2022 Mar 28];23(21):2947–2948. https://pubmed.ncbi.nlm.nih.gov/17846036/. doi:10.1093/BIOINFORMATICS/BTM404

21. Ma J, Pan Z, Huang J, Sun M, Lu C, Yao H. The Hcp proteins fused with diverse extended-toxin domains represent a novel pattern of antibacterial effectors in type VI secretion systems. Virulence. 2017;8(7):1189–1202. 10.1080/21505594.2017.1279374. doi:10.1080/21505594.2017.1279374

22. Ma J, Sun M, Dong W, Pan Z, Lu C, Yao H. PAAR-Rhs proteins harbor various C-terminal toxins to diversify the antibacterial pathways of type VI secretion systems. Environmental Microbiology. 2017 [accessed 2019 May 21];19(1):345–360. http://www.ncbi.nlm.nih.gov/pubmed/27871130. doi:10.1111/1462-2920.13621

23. Marchler-Bauer A, Lu S, Anderson JB, Chitsaz F, Derbyshire MK, DeWeese-Scott C, Fong JH, Geer LY, Geer RC, Gonzales NR, Gwadz M. Hurwitz DI, Jackson JD. Ke Z, Lanczycki CJ, Lu F, Marchler GH, Mullokandov M, Omelchenko MV, Robertson CL, Song JS, Thanki N, Yamashita RA, Zhang D, Zhang N, Zheng C, Bryant SH. CDD: A Conserved Domain Database for the functional annotation of proteins. Nucleic Acids Research. 2011 [accessed 2020 Nov 23];39(SUPPL. 1). https://pubmed.ncbi.nlm.nih.gov/21109532/. doi:10.1093/nar/gkq1189

24. Martorelli L, Albanese A, Vilte D, Cantet R, Bentancor A, Zolezzi G, Chinen I, Ibarra C, Rivas M, Mercado EC, Cataldi A. Shiga toxin-producing Escherichia coli (STEC) O22:H8 isolated from cattle reduces E. coli O157:H7 adherence in vitro and in vivo. Veterinary Microbiology. 2017;208(June):8–17. doi:10.1016/j.vetmic.2017.06.021

25. Navarro-Garcia F, Ruiz-Perez F, Cataldi Á, Larzábal M. Type VI secretion system in pathogenic escherichia coli: Structure, role in virulence, and acquisition. Frontiers in Microbiology. 2019;10(AUG):1–17. doi:10.3389/fmicb.2019.01965

26. Nguyen LT, Schmidt HA, Von Haeseler A, Minh BQ. IQ-TREE: a fast and effective stochastic algorithm for estimating maximum-likelihood phylogenies. Molecular biology and evolution. 2015 [accessed 2022 Mar 28];32(1):268–274. https://pubmed.ncbi.nlm.nih.gov/25371430/. doi:10.1093/MOLBEV/MSU300

27. Okuyama M, Mori H, Chiba S, Kimura A. Overexpression and characterization of two unknown proteins, YicI and YihQ, originated from Escherichia coli. Protein Expression and Purification. 2004 [accessed 2020 Nov 6];37(1):170–179. https://pubmed.ncbi.nlm.nih.gov/15294295/. doi:10.1016/j.pep.2004.05.008

28. Répérant M, Porcheron G, Rouquet G, Gilot P. The yicJI metabolic operon of Escherichia coli is involved in bacterial fitness. FEMS Microbiology Letters. 2011;319(2):180–186. doi:10.1111/j.1574-6968.2011.02281.x

29. Rivas M, Miliwebsky E, Chinen I, Roldán CD, Balbi L, García B, Fiorilli G, Sosa-Estani S, Kincaid J, Rangel J, Griffin, P. M. Chillemi G, Baschkier A, Manfredi E, Ojea J, Caletti MG, Amoedo D, Martin S, Vallès P, Lo Giudice P, Pesle S, Principi I, Peralta M, Marsano De Mollar MC, Hoekstra RM. Characterization and epidemiologic subtyping of Shiga toxin-producing Escherichia coli strains isolated from hemolytic uremic syndrome and diarrhea cases in Argentina. Foodborne Pathogens and Disease. 2006 [accessed 2020 Nov 6];3(1):88–96. https://pubmed.ncbi.nlm.nih.gov/16602984/. doi:10.1089/fpd.2006.3.88

30. Russell AB, Leroux M, Hathazi K, Agnello DM, Ishikawa T, Wiggins PA, Wai SN, Mougous JD. Diverse type VI secretion phospholipases are functionally plastic antibacterial effectors. Nature. 2013 [accessed 2020 Nov 6];496(7446):508–512. https://www.nature.com/articles/nature12074. doi:10.1038/nature12074

31. Russell AB, LeRoux M, Hathazi K, Agnello DM, Ishikawa T, Wiggins PA, Wai SN, Mougous JD. Diverse type VI secretion phospholipases are functionally plastic antibacterial effectors. Nature. 2013 [accessed 2016 Apr 9];496(7446):508–12. http://www.ncbi.nlm.nih.gov/pubmed/23552891. doi:10.1038/nature12074

32. Russell AB, Peterson SB, Mougous JD. Type VI secretion system effectors: Poisons with a purpose. Nature Reviews Microbiology. 2014 [accessed 2020 Nov 6];12(2):137–148. https://pubmed.ncbi.nlm.nih.gov/24384601/. doi:10.1038/nrmicro3185

33. Santos MNM, Cho ST, Wu CF, Chang CJ, Kuo CH, Lai EM. Redundancy and Specificity of Type VI Secretion vgrG Loci in Antibacterial Activity of Agrobacterium tumefaciens 1D1609 Strain. Frontiers in Microbiology. 2020;10(January):1–15. doi:10.3389/fmicb.2019.03004

34. Ting SY, Bosch DE, Mangiameli SM, Radey MC, Huang S, Park YJ, Kelly KA, Filip SK, Goo YA, Eng JK, Allaire M, Veesler D, Wiggins PA, Peterson S, Brook M, Joseph D. Bifunctional Immunity Proteins Protect Bacteria against FtsZ-Targeting ADP-Ribosylating Toxins. Cell. 2018;175(5):1380-1392.e14. 10.1016/j.cell.2018.09.037. doi:10.1016/j.cell.2018.09.037

35. Vazquez-Lopez J, Navarro-Garcia F. In silico Analyses of Core Proteins and Putative Effector and Immunity Proteins for T6SS in Enterohemorrhagic E. coli. Frontiers in Cellular and Infection Microbiology. 2020;10(May):1–13. doi:10.3389/fcimb.2020.00195

36. Wan B, Zhang Q, Ni J, Li S, Wen D, Li J, Xiao H, He P, Ou H, Tao J, Teng Q, Lu J, Wu W, Yao YF. Type VI secretion system contributes to Enterohemorrhagic Escherichia coli virulence by secreting catalase against host reactive oxygen species (ROS). Zamboni DS, editor. PLOS Pathogens. 2017 [accessed 2020 Nov 6];13(3):e1006246. https://dx.plos.org/10.1371/journal.ppat.1006246. doi:10.1371/journal.ppat.1006246

37. Whitney JC, Chou S, Russell AB, Biboy J, Gardiner TE, Ferrin MA, Brittnacher M, Vollmer W, Mougous JD. Identification, structure, and function of a novel type VI secretion peptidoglycan glycoside hydrolase effector-immunity pair. Journal of Biological Chemistry. 2013 [accessed 2020 Nov 6];288(37):26616–26624. /pmc/articles/PMC3772208/?report=abstract. doi:10.1074/jbc.M113.488320

38. Zhang D, Iyer LM, Aravind L. A novel immunity system for bacterial nucleic acid degrading toxins and its recruitment in various eukaryotic and DNA viral systems. Nucleic Acids Research. 2011;39(11):4532–4552. doi:10.1093/nar/gkr036

39. Zimmermann L, Stephens A, Nam SZ, Rau D, Kübler J, Lozajic M, Gabler F, Söding J, Lupas AN, Alva V. A Completely Reimplemented MPI Bioinformatics Toolkit with a New HHpred Server at its Core. Journal of Molecular Biology. 2018;430(15):2237–2243. doi:10.1016/j.jmb.2017.12.007

